# Rapid, multianalyte detection of opioid metabolites in wastewater

**DOI:** 10.1101/2021.09.09.459680

**Authors:** Narendra Kumar, Muhit Rana, Michael Geiwitz, Niazul Islam Khan, Matthew Catalano, Juan C. Ortiz-Marquez, Hikari Kitadai, Andrew Weber, Badawi Dweik, Xi Ling, Tim van Opijnen, Avni Argun, Kenneth S. Burch

## Abstract

By monitoring opioid metabolites, wastewater-based epidemiology (WBE) could be an excellent tool for real-time information on consumption of illicit drugs. A key limitation of WBE is the reliance on costly laboratory-based techniques that require substantial infrastructure and trained personnel, resulting in long turnaround times. Here, we present an aptamer-based graphene field effect transistor (AptG-FET) platform for simultaneous detection of three different opioid metabolites. This platform provides a reliable, rapid, and inexpensive method for quantitative analysis of opioid metabolites in wastewater (WW). The platform delivers a limit of detection (LOD) 2-3 orders of magnitude lower than previous reports, but in line with the concentrations range (pg/ml to ng/ml) of these opioid metabolites present in real samples. To enable multianalyte detection we developed a facile, reproducible, and high yield fabrication process producing twenty G-FETs with integrated side gate platinum (Pt) electrodes on a single chip. Our devices achieved the simultaneous and selective multianalyte detection of three different metabolites: Noroxycodone (NX), 2-ethylidene-1,5-dimethyl-3,3-diphenylpyrrolidine (EDDP), and Norfentanyl (NF) in wastewater.

## 1. Introduction

Effective responses to the opioid epidemic require real-time and local information on the type and usage frequency of illicit drugs.^1,2^ A new approach has emerged that avoids many of the difficulties associated with individual testing, namely wastewater-based epidemiology (WBE). This strategy enables community tracing of drug metabolites and tracking the spread of infectious diseases.^3–7^ Wastewater monitoring can provide near real-time feedback on the introduction and continued usage of psychoactive substances without stigmatizing communities, households or individuals.^8,9^ However, successful WBE requires a highly sensitive and specific detection technique as the concentrations of metabolites in wastewater are very low (pg/ml to ng/ml) due to excessive dilution.^10,11^ The current gold standard method to detect these drug metabolites is high-pressure liquid chromatography tandem mass spectrometry (HPLC-MS) which requires advanced equipment, sample analyses, skilled personnel, and cannot be performed on site.^12–14^ As such, the implementation of HPLC-MS has been limited by a long turnaround time and cost. Therefore, to make WBE more meaningful and reliable, a rapid, highly sensitive, cost effective, and easy to use detection method is required for on-site analysis of drug metabolites.

Attempts to achieve these aims have employed different techniques such as colorimetric, fluorescence, surface-enhanced Raman spectroscopy (SERS), lateral flow immunoassay (LFIA), and electrochemical detection.^10,15–21^ In these approaches, optical detection using nanomaterials-based aptasensors have been extensively investigated for rapid analysis of illicit drugs.^15^ Nanomaterials are utilized to achieve high sensitivity and lower limit of detection (LOD) values while aptamer probes possess excellent affinity, stability at room temperature, smaller size, and can be chemically synthesized on a large scale and at low-cost.^16,22^ For example, a gold nanoparticles conjugated assay was reported with a LOD of 0.5 nM (0.15 ng/ml) and 3.3 nM (1 ng/ml) for Methadone and Cocaine respectively.^23^ The optical assays-based techniques using nanomaterials are limited by high LOD, miniaturization, complex equipment, and cost. On the other hand, the LFIA and electrochemical sensors have the capability to solve several challenges, but have yet to achieve high sensitivity and stability in real wastewater samples.^17,18^ For instance, a nafion-coated carbon nanotube electrode can specifically detect Oxycodone with a LOD of 85 nM (27 ng/ml), which is quite high considering the very low amount (∼pg/ml) for several drug metabolites present in wastewater samples.^10,19,24,25^ Similarly, an LFIA based sensor showed sensitivity (LOD) values of 5-50 ng/ml for detecting Fentanyl (Norfentanyl as metabolite) but were only tested with urine, PBS, and saliva samples and not in wastewater.^18^ The LFIA still suffers from low sensitivity and quantification while the electrochemical sensors require complex fabrication due to dependence on nanomaterial modification to achieve the desired detection limit.^26,27^ Moreover, most experimental approaches such as enzyme-linked immunosorbent assay (ELISA) and LFIA mostly rely on antibodies that might be incompatible with waster while suffering from inconsistencies between vendors and product lots.^20,21^ To the best of our knowledge, graphene field effect transistors (G-FETs) with aptamer probes have not yet been implemented into a field deployable wastewater sensor. Herein we report the development of a miniaturized G-FET platform utilizing highly specific aptamers (AptG-FET) (Figure 1) for rapid, sensitive, and simultaneous detection of drug metabolites in wastewater.

**Figure 1.**
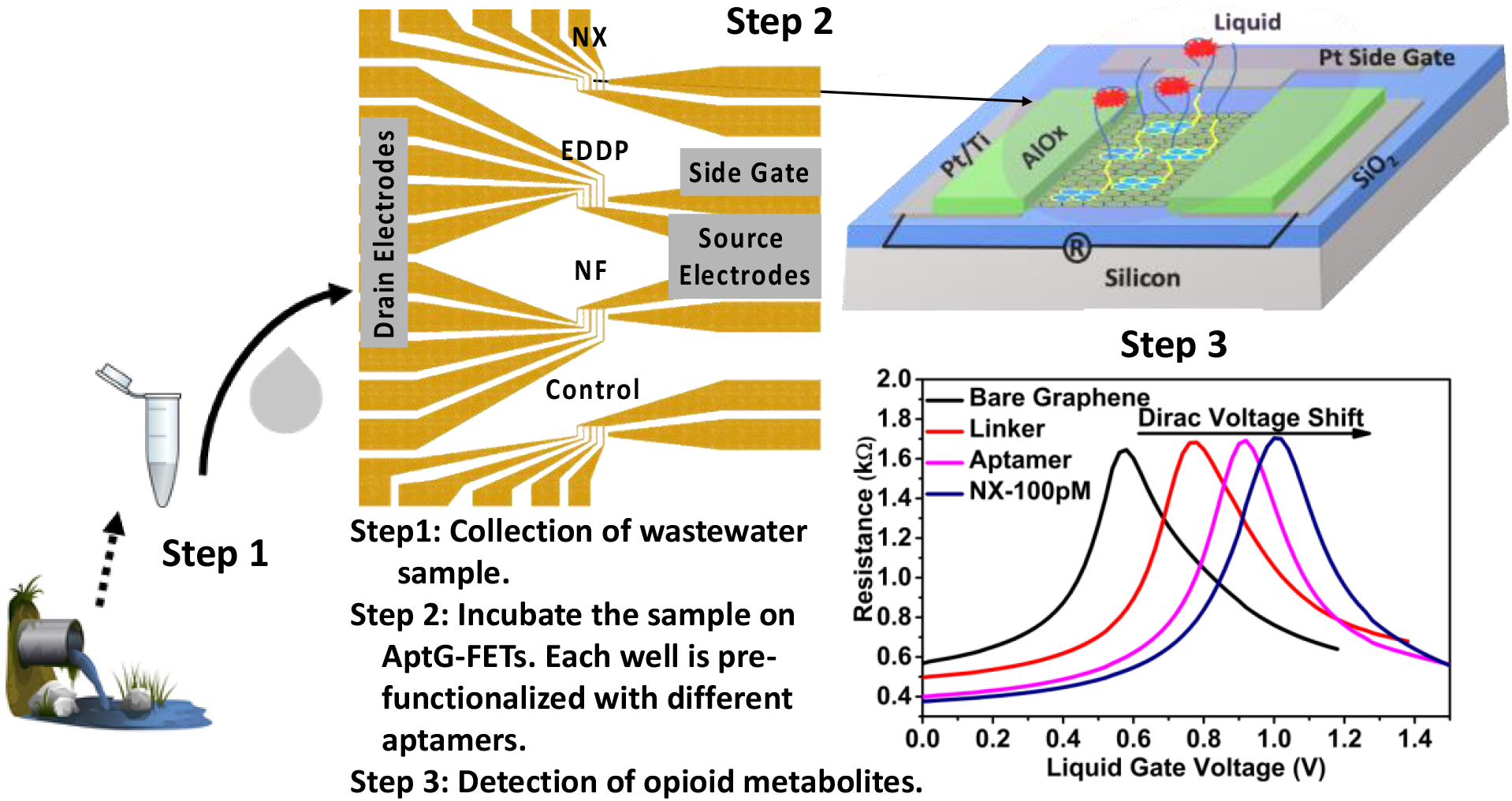
Schematic illustration of an onsite chip-based rapid detection platform for near real-time monitoring of opioid metabolites in wastewater using AptG-FET sensor technology. Step 1, wastewater collection, filtration, and dilution (as needed); Step 2, on chip sample (10 μL) incubation; Step 3, sensor characterization of the sensor to estimate the concentration of targeted drug metabolites.

In the last decade, G-FET based biosensors have emerged as sensors with a large potential due to their high sensitivity, biocompatibility, non-covalent functionalization, and scalable fabrication on various substrates.^28–33^ The electrical resistance of graphene is highly sensitive to the target bio-analytes (or the conformal changes of the probe), enabling direct and rapid readout.^28,30,34,35,36,37^ In our recent work, we demonstrated a highly sensitive G-FET for the detection of biomarkers such as CA-I (oral diseases biomarker) in saliva,^38^ and antibiotic resistant bacteria, both at clinically relevant concentrations.^39^ However, the G-FET design in our prior work was limited to detection of a single target, with each chip functionalized with a single probe, provided minimal passivation, and required a platinum (Pt) wire as a separate reference electrode.

Thus, our previous G-FETs were not appropriate for WBE, where one requires multi-analyte detection and robust devices on a self-contained chip. To achieve this, we developed a facile and reproducible fabrication process for multiple, isolated sets of G-FETs with a platinum (Pt) reference electrode on a single chip (1.2 cm × 1.2 cm). We segregated the chip in four different sets of devices and Polydimethylsiloxane (PDMS) wells were mounted to functionalize the chip with four different probes for multianalyte detection of opioid metabolites (Figure 1). To further ensure the accuracy and robustness of the sensor, we kept five G-FET devices in each well to average the output signal and calculate the standard deviation. Furthermore, we eliminated the external Pt reference electrode by fabricating on chip side gate electrodes. To ensure the robustness of the G-FETs in wastewater we passivated the devices with an aluminum oxide (AlOx) layer, exposing only the active area of graphene for selected attachment of probes. To keep the graphene surface pristine, a new process protocol was developed that is easy to fabricate and ensures excellent stability (see SI 1.1). The devices were made in our unique cleanroom in a glovebox, significantly reducing the cost of production and fabrication time as well as limiting the exposure of the graphene to atmospheric contaminants.^40^

The design and fabrication enhancements enable our new AptG-FET platform to simultaneously detect four different drug metabolites from a single sample of wastewater. To demonstrate this, we first validated the aptamer’s binding affinity with the respective drug metabolites in standard buffer and wastewater using plasmonic and electrochemical detection techniques. Then, one of the aptamers was functionalized over the G-FET sensor to confirm the sensitivity, selectivity, and detection limit. Finally, the multianalyte detection of all three targeted metabolites were performed on the same chip and their sensitivity, affinity, and selectivity were tested.

## 2. Results and Discussion

### 2.1 Selection and Validation of Aptamers

Three different opioid metabolites were chosen in this study, namely Noroxycodone (NX), 2-ethylidene-1,5-dimethyl-3,3-diphenylpyrrolidine (EDDP) and Norfentanyl (NF). NX is commonly used for analgesic effects and pain management, EDDP is prescribed to overcome opioid addiction, and fentanyl is a more potent, synthetic opioid used in intravenous anesthetics.^41–43^ To ensure the selected aptamers have strong binding affinities, we performed a standard nanoplasmonic assay for characterization of aptamers binding to their specific targets. We focused on confirming the binding time, sensitivity, specificity, and dynamic detection range.

Briefly, citrate-reduced gold nanoparticles (AuNP) possess negative charge and the electrostatic repulsive forces account for their characteristic red color when dispersed in a medium.^15,23^ In the presence of a negatively charged aptamer and ∼30 mM NaCl, a negative charge cloud protects AuNPs from any aggregation (Figure 2a). When an aptamer binds to the target, it leaves the particle surface, reducing the inter-particle distance. Next salt-induced aggregation of the AuNP takes place, resulting in red-to-purple color transition in less than a minute.^44–46^ This simple mechanism allows us to obtain quantitative binding information by monitoring the optical density (OD) at each wavelength and comparing the ratio of (OD_520_/OD_700_) (Figure 2a inset), demonstrating aptamer functionality in buffer and challenging wastewater matrices. A strong peak in OD_520_ is observed in absence of the target which upon target binding is reduced by ∼40%. We have successfully used this strategy to validate aptamers for NX, NF and EDDP in buffer. This assay is also used to determine LOD (≥ 6.05 nM) and selectivity of each aptamer in the relevant wastewater media for individual target drug metabolites.

**Figure 2.**
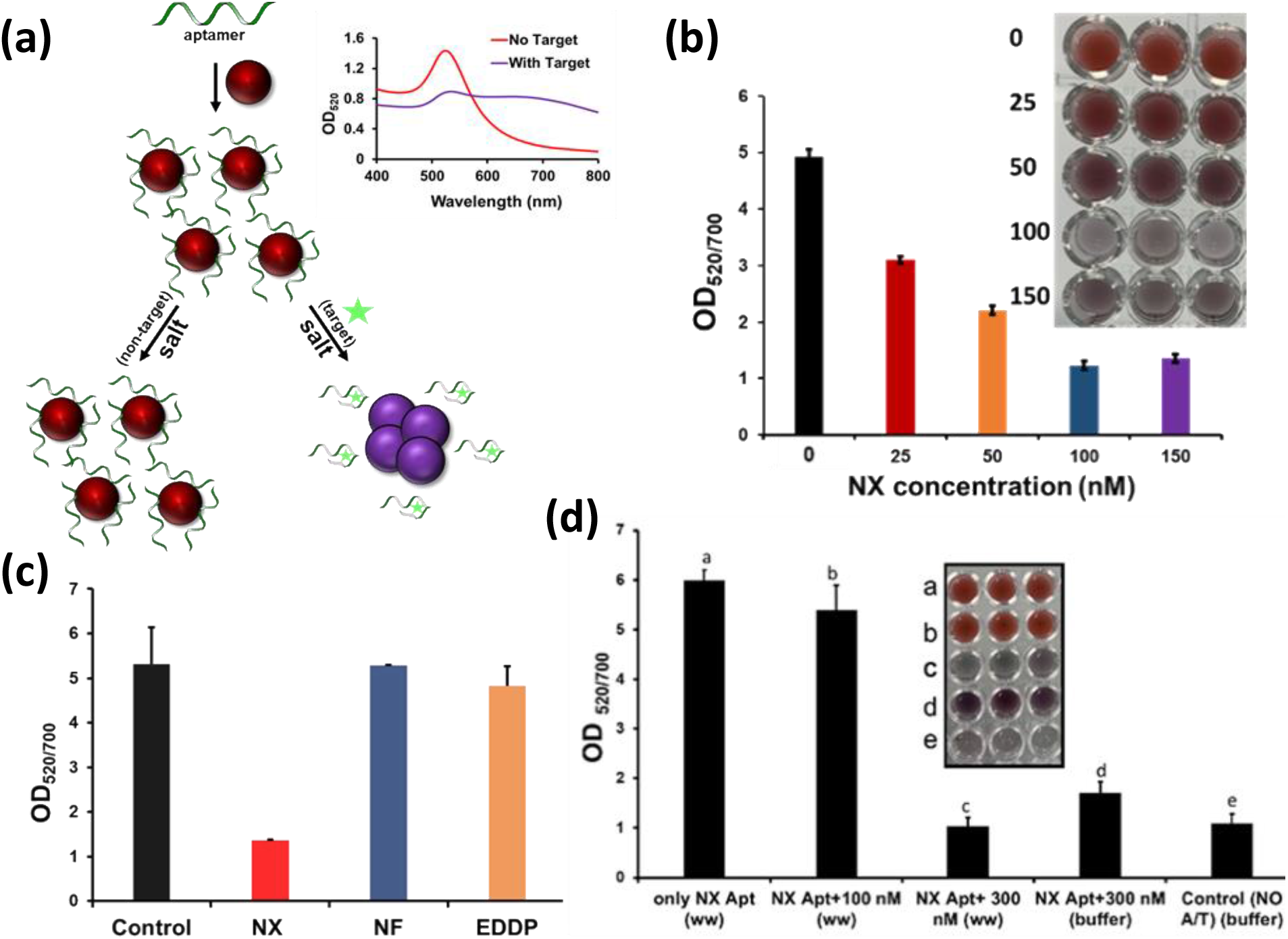
Aptamer validation procedure that utilizes the relative optical density change resulting from aggregation of AuNPs in presence of targets. (a) A schematic shows the change in color of AuNPs with and without presence of target analyte (inset change in OD). (b) Characterization of AuNPs functionalized with an opioid aptamer (such as-NX) to obtain the sensitivity by measuring change in OD values. (c) Specificity analysis of NX-aptamer in buffer (1x PBS), very little variation observed with EDDP and NF while a strong decrease in OD with NX target. (d) NX detection in filtered and diluted wastewater, a minimum of 300 nm NX target required against 100 nM aptamer to achieve significant shift in signal (inset change in OD). All experiments were performed in triplicate (n=3).

Using the colorimetric assay, we performed sensitivity and specificity analysis of the aptamers for NX, EDDP and NF in 1x PBS (phosphate buffer saline), pH 7.4. Figure 2b demonstrates a dose-dependent linear correlation between the absorbance reading (OD_520_/OD_700_) and various NX (one of the representative opioids in our study) levels with a LOD of 6.05 nM (1.8 ng/ml). As expected, a more drastic color change was observed when higher dose of NX was introduced in buffer. We have observed similar findings for other opioid aptamers (EDDP and NF) with their target drug analytes.

To determine the specificity of these aptamers, we tested them with non-complementary target analytes. As shown in Figure 2c, in the presence of NX aptamer, when a non-complementary target analyte (NF or EDDP) was introduced, the absorbance reading showed nearly no difference compared to the control (only aptamer) result. The salt-induced aggregation only takes place with the correct aptamer and NX target, and the ratio (OD_520_/OD_700_) equivalent to the aggregation state is measured. These data confirm the absence of any false positive or false negative detection and thus verified the specificity of the chosen aptamer (high rates of false positives and false negatives are major drawbacks for existing immunoassays that rely on antibody binding).^20,21^

Having established the sensitivity and specificity of the aptamers in buffer solutions, we turned to confirm the aptamers would work with wastewater samples. The influent (raw and untreated) wastewater samples were collected from The Massachusetts Alternative Septic System Test Center (MASSTC), located in Sandwich, MA. To avoid interference in detection due to the presence of the bigger objects and species, filtering as received wastewater sample was performed using a 0.22-micron filter.^14^ To achieve the successful binding of aptamers, the sample was further diluted to 1:20 in binding buffer solution (1x PBS +2 mM MgCl_2_ +1% Methanol) and spiked with different concentrations of opioid metabolites. As seen in Figure 2d, a distinguishable detection of opioids in wastewater samples via the AuNP requires at least 300 nM (90 ng/ml) of NX target compared to 100 nM of NX target when measured in buffer. So, the working condition of the aptamer: target is 1:1 (Figure 2c) in buffer and 1:3 (Figure 2d) in wastewater samples. The increase of LOD in wastewater samples is attributed to the interference caused by the presence of several other analytes and species. Overall, this data indicates the functionality and selectivity of all aptamers for detection of the three opioid metabolites (NX, NF, and EDDP) both in buffer and in wastewater samples (NF and EDDP shown in Figure S3) These findings were further validated with electrochemical impedance spectroscopy (EIS) using screen printed electrodes in 1x PBS which showed an LOD of 10 nM (3 ng/ml) for NX target (see SI 1.3). However, no sensitive detection of opioid metabolites was observed in wastewater samples when tested with EIS.

### 2.2. Detection of NX with G-FET in 1x PBS

Having confirmed the aptamer’s binding and specificity utilizing nano-plasmonic and electrochemical detection, we turned to our G-FET sensing platform. There are two different ways to operate a G-FET to perform biosensing; one is back gating through silicon and another through an ionic liquid. Traditional back-gated G-FETs offer reference electrode free devices but require substantial voltages (>60V) with special electronics.^30^ Liquid gated G-FET sensors significantly lower the required voltage (below 2 V) as well as keep the probes and analytes in their original size and conformation. Our prior work, as well as others^38,39^, show liquid gating is a reliable approach with less complex electronics required for backgated FET. In this work, we utilized our upgraded G-FET platform having on-chip coplanar Pt side gate electrodes which provides a miniaturized G-FET platform and allows upscaling of the number of devices on the same chip while measuring them simultaneously (see Figure 1). A detailed description of the multiplexed G-FET design is provided in the following section, with a schematic shown in Figure 3a along with a microscopic image of the device in Figure 3b. Furthermore, to achieve selective functionalization in the channel region, a graphene sensing window of 10 μm × 40 μm was defined by depositing and selectively etching 50 nm of AlO_x_. This thickness of AlO_x_ around the contact pads further improves the stability of the sensor by passivating the source/drain electrodes and minimizing leakage current. For the specific detection of the NX target, the G-FETs were first functionalized with 1mM 1-pyrenebutyric acid N-hydroxysuccinimide ester (PBASE) linker dissolved in dimethylformamide (DMF) for 1 h and rinsed with DMF, isopropyl alcohol (IPA), and DI water. Then, NX-aptamer with 10 μM concentration was incubated for 1 h and rinsed with PBS, and DI water. Raman spectroscopy confirmed the attachment of aptamers to graphene (see Figure S5). To obtain a resistance vs liquid gate voltage plot, the measurements were performed in 0.01x PBS to minimize the Debye screening effect.^38^ Figure 3c shows the shift in Dirac voltage (*V*_*D*_) after successive functionalization with PBASE linker and NX-aptamers. Further shift in *V*_*D*_ was observed upon incubation of NX target at concentrations of 1 nM and 10 nM while saturating when measured with 100 nM. Several devices have been tested to obtain the concentration dependent shift in *V*_*D*_ as shown in Figure 3d. No significant shift was observed at 1pM while an incremental shift was recorded at 10 pM and higher values of NX concentration (Figure S6).

**Figure 3.**
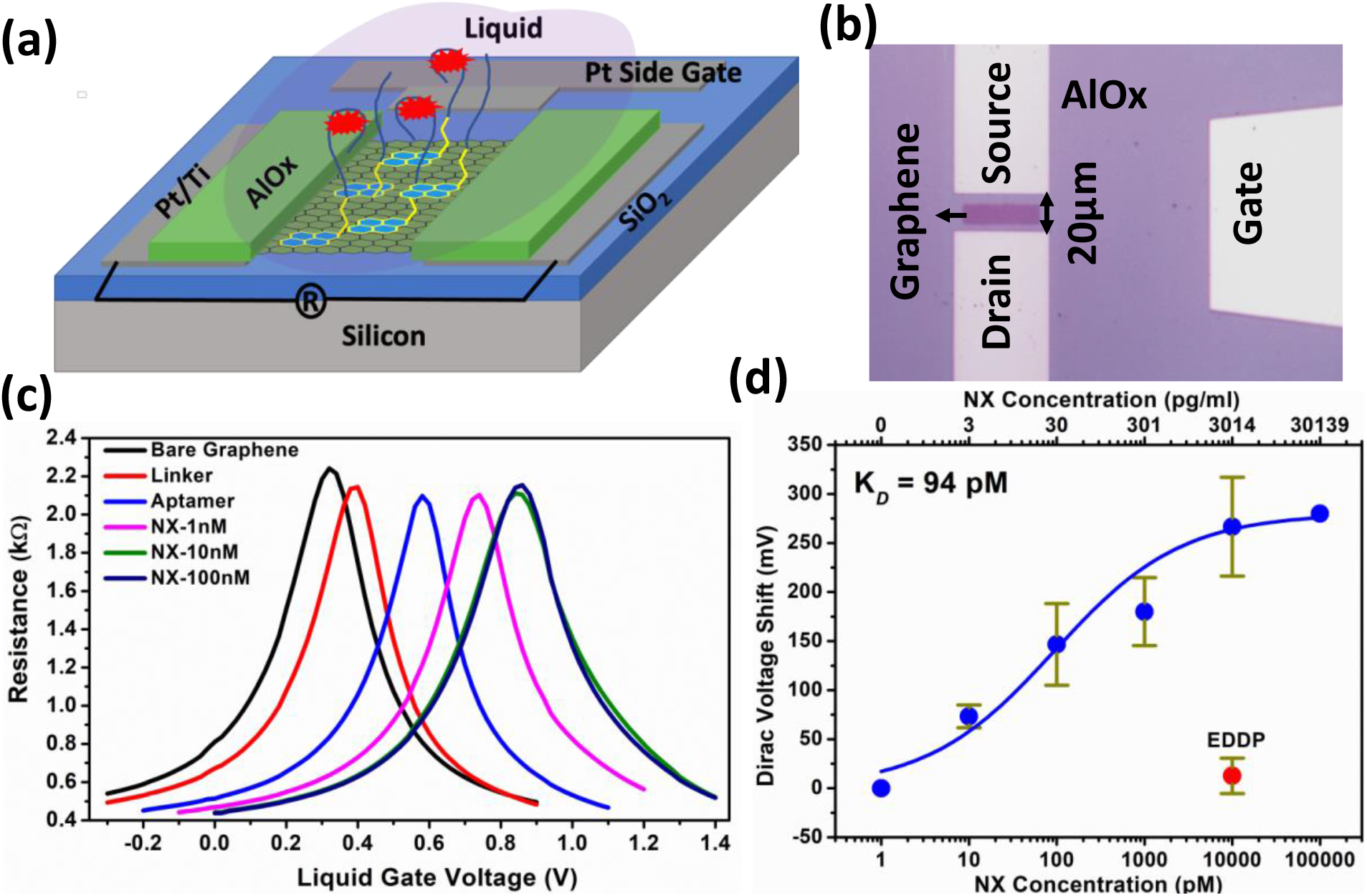
(a) Schematic of fabricated G-FET (b) Microscopic image of single G-FET with source/drain and side gate electrode, graphene sensing window with AlO_x_ passivation (c) G-FET characteristics upon functionalization with aptamer probes and detection of Noroxycodone (d) Concentration dependence of Dirac voltage shift along with shift from high concentration of negative control (EDDP). Error bars were calculated with the data from 3-5 devices at each concentration.

To understand the binding kinetics of the aptamers with NX, we fit the calibration curve to Hill’s equation: 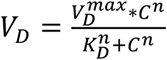 Here *V*_*D*_ is the measured Dirac voltage shift at different concentrations of 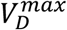 is the Dirac voltage shift when all the binding sites are saturated, C is the concentration of NX, *K*_*D*_ is the dissociation constant, and n is the Hill’s coefficient. As shown in Figure 3d, the resulting fit provides an excellent description of the concentration dependence. Interestingly, we find a *K*_*D*_ of 94 pM which is much better than the conventional fluorescent and HPLC techniques yet comparable to G-FET based biosensors used for other types of biomarkers.^19,23,26,47^ From the fits we found the LOD was approximately 10 pM which is nearly two orders of magnitude higher than that obtained with fluorescent assay and mass spectrometry^19,48^ and is comparable to that obtained for oxycodone utilizing electrodes modified with different complex nanostructures and immobilization process.^26,27^ To confirm the specificity, we tested the NX-aptamers functionalized G-FET with 100 nM EDDP, only a small shift of 8 mV was seen which is much lower than that (∼60 mV) obtained with 10 pM of NX concentration (see Figure 3d). This confirmed that aptamers used for NX detection are also highly specific.

### 2.3. Simultaneous detection of three different targets in wastewater

After confirming the sensitivity and specificity of the NX-aptamer in 1x PBS, we moved to test the three different opioid metabolites (NX, EDDP, and NF) together in real wastewater samples. To test them all together, we utilized our upgraded G-FET detection platform. We note the entire platform is extremely small enabling ease of use and portability (Figure 4a). A microscopic image of one well showing four devices and a side gate electrode is shown in Figure 3b. Each PDMS well can hold 10 µL of solution. The devices in all the wells are measured simultaneously before and after functionalization. Three different wells were first functionalized with PBASE linker followed by three different aptamers, i.e. NX-Apt, EDDP-Apt, and NF-Apt while the fourth well is used as a control. In this way, each well has an aptamer for one respective target. The pre-functionalized G-FETs were tested with raw wastewater which resulted in no significant variation in characteristics (i.e. - *V*_*D*_, mobility, resistance) confirming the stability of the devices. However, no shift in *V*_*D*_ was observed until 1μM of NX target because of the interference caused by several other analytes and species. So, we utilized the filtered and diluted (20x in binding buffer) wastewater spiked with different concentration of targets. Then, the simultaneous detection of all three targets was performed. Representative plots of resistance vs voltage at each stage of functionalization and detection of different concentrations of targets are shown in Figure S7. To avoid non-specific binding at the graphene surface and minimize the interference of different analytes likely present in wastewater, both end amine-terminated polyethylene glycol (PEG 1 KDa) was mixed in a 1:1 ratio with the aptamers during functionalization.^29^ This resulted in more stable behavior with minimized drift and standard deviation in calculated error bars (see data without PEG in Figure S8). The other two targets were utilized as negative controls while detecting the third target. For example, high concentrations (100 nM) of EDDP and NF were used as negative controls for the well functionalized with NX-Apt, then successive detection of different NX concentrations was performed (Figure 4b). No significant shift in *V*_*D*_ was observed with EDDP and NF while an incremental voltage shift was observed with increasing NX concentration. The shift in *V*_*D*_ saturated around 100 nM suggesting most available binding sites of aptamer probes are occupied with added integration of target analyte. Similar trends of concentration dependent voltage shifts were observed with other the two targets (EDDP and NF) as shown in Figure 4c and 4d. Juxtaposed to NX and EDDP, the devices functionalized with NF-Apt showed a higher shift (∼50 mV) with NF target concertation at 10 pM, which could be attributed to the higher binding affinity of this aptamer (Figure 4d). However, the devices functionalized with NF-Apt also showed some level of non-specificity to NX and EDDP targets of concentrations (100 nM) with a voltage shift of ∼40 mV (Figure 4d).

**Figure 4.**
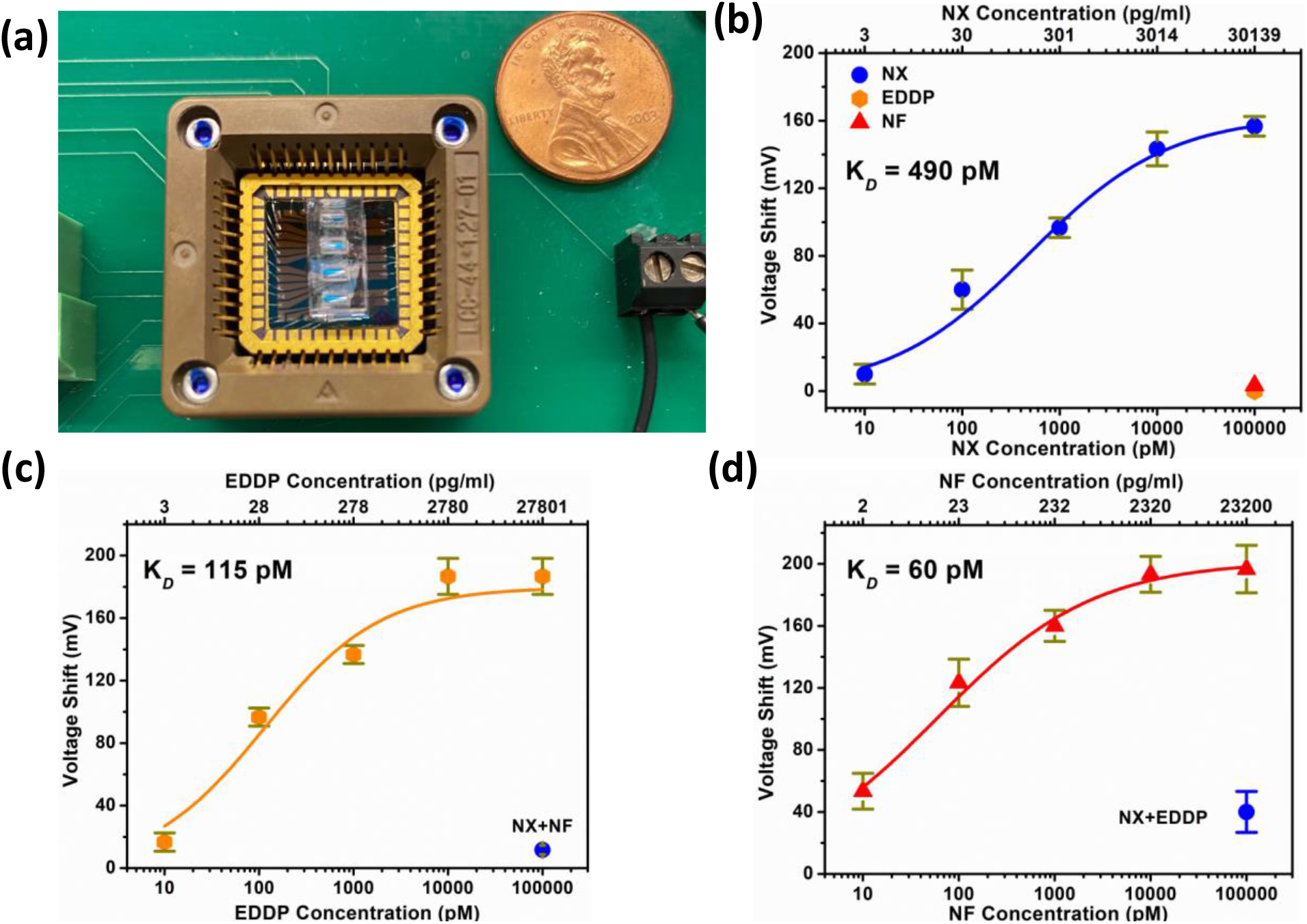
(a) G-FET chip with 4 PDMS wells (b) Calibration curve for NX in wastewater with negative control of EDDP and NF, K_D_ value of 490 pM and LOD of 126 pM (c) Calibration curve for EDDP with negative controls of NX and NF, K_D_ value of 115 pM and LOD of 96 pM (d) Calibration curve for NF with negative controls of NX and EDDP, K_D_ value of 60 pM and LOD of 183 pM. Error bars were calculated with the data from 5 devices at each concentrations.

To confirm the binding kinetics of these aptamers and targets in wastewater, the calibration curves obtained with all three AptG-FETs were fitted to the Hill’s equation. They all showed strong binding affinity as evident from the resulted K_D_ values of 490 pM, 115 pM, and 60 pM for NX, EDDP, and NF, respectively. The higher binding affinity (K_D_) of aptamers and sensitivity of G-FETs resulted in significantly lower statistical LOD values of 126 pM (38 pg/ml), 96 pM (27 pg/ml), and 183 pM (42 pg/ml) for NX, EDDP, and NF respectively. These obtained values of K_D_ and LOD are 2 to 3 orders of magnitude lower than those reported in the literature for all three drug metabolites.^1,12,15,19,23^ Considering the 1:20 dilutions of used wastewater, these obtained LOD values are still on the order of 100 pg/ml with the detection range of pg/ml to ng/ml which is in line with the concentration range of metabolites present in wastewater.^10,11,24,25^

### 2.4 Selective detection in wastewater

After confirming the sensitivity and specificity of the opioid metabolites in wastewater, their selectivity was tested. Specifically, the one specific target was mixed with other two non-specific targets and detection was performed with G-FETs to see the hinderance in signal in comparison with that detected individually. Figure 5a shows the signal obtained from two different concentrations of NX target mixed with similar amounts of EDDP and NF, obtained values are slightly lower than that tested with NX alone as a target. A similar test was also performed for EDDP which shows the voltage shift much closer to that obtained with a single target present (Figure 5b). However, the voltage shift with NF when mixed with NX and EDDP was significantly lower as compared to the other two targets (Figure 5c). This lower shift could be attributed to the interference of other targets as we have already observed some level of unspecific binding with NF-aptamers. This confirms that these AptG-FETs possess promising selectivity level along with their high sensitivity and affinity in wastewater.

**Figure 5.**
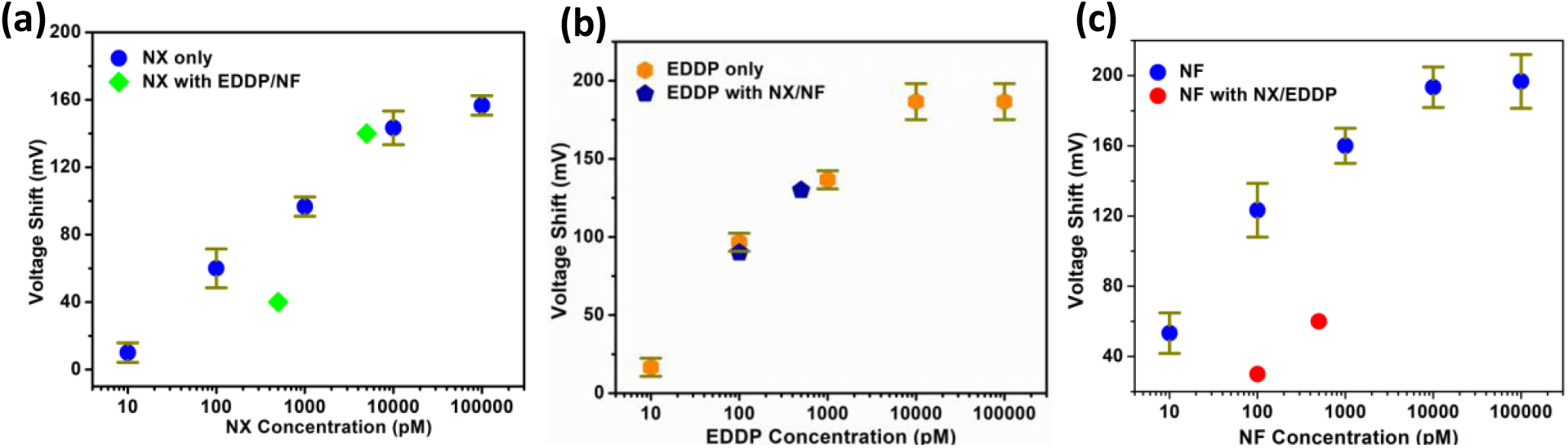
(a) Selective detection of NX from mixed samples with EDDP and NF, (b) Selective detection of EDDP from mixed samples with NX and NF, (c) Selective detection of NF from mixed samples with NX and EDDP.

## 3. Conclusion and future directions

In summary, we demonstrated the capabilities of aptameric G-FET sensors for rapid, selective, and simultaneous detection of three different drug metabolites in wastewater. Our AptG-FET platform provides multianalyte detection on a single chip (1.2 cm × 1.2 cm) which consists of four different PDMS wells each having five devices, on chip coplanar side gate electrodes, and passivation layer of AlO_x_ layer. Our presented AptG-FET platform showed high specificity, sensitivity, and selectivity for all three opioid metabolites used in this work. The achieved LOD values of 38, 27, and 42 pg/ml for NX, EDDP, and NF respectively, are well in line with the desired limits in wastewater and are 2-3 orders of magnitude better that what has been achieved with other techniques.^10,11,18,19,23^ All AptG-FETs have shown high binding affinity with K_D_ values of 490 pM, 115 pM, and 60 pM for NX, EDDP, and NF respectively. Thus, the presented AptG-FET platform is capable to be utilized for real time monitoring of illicit drugs in wastewater and can provide a boost to the WBE. Our current device design is straight forward to scale to a larger number of wells for detecting an array of analytes. In the future, the same platform, with different probes, could be utilized for wastewater-based monitoring of a variety of analyte types including pathogens and other disease biomarkers in local health monitoring and epidemiology studies. Furthermore, our device’s design, size, rapid response, multianalyte capabilities, scalability and ease of operation enable a new era of wastewater epidemiology at the local level.

## 4. Materials and methods

### 4.1. Materials

1-pyrenebutyric acid N-hydroxysuccinimide ester (PBASE) linker and Dimethylformamide (DMF) were obtained from Sigma Aldrich. All aptamers (5’-amine-Aptamer-3’: Norfentanyl: CFA0071-GP5-25 AKA-H6AAZ; NX: CFA0079-GP5-25, AKA-H4LFD and EDDP: CFA0661-GP5-25) and their resuspension buffer were purchased from Base Pair which has developed aptamers that are capable of binding noroxycodone, EDDP, and norfentanyl, and has readily available aptamers for Morphine. Target noroxycodone hydrochloride, EDDP, and norfentanyl oxalate were purchased as ampules of 1 mL with concentration 1mg/mL in methanol (as free base) from Sigma Aldrich, St. Louis, MO 63103, USA. Disposable screen-printed carbon electrodes (SPCEs) were purchased from Metrohm (DRP-110CNT) with carbon working and auxiliary electrodes and silver as the reference electrode where the working electrodes were modified with carboxyl functionalized with multi-walled carbon nanotubes (MWCNT-COOH). Ambion™ DEPC-treated nuclease-free water (0.2 µm filtered and autoclaved) was purchased from Invitrogen, Thermo Fisher Scientific (Waltham, MA, USA) and utilized in all studies. To avoid any DNase contamination, DNA Away (DNA Surface Decontaminant) was purchased from Thermo Scientific and used before performing any experiment. All other reagents and buffers were purchased from Sigma-Aldrich, St. Louis, MO 63103, USA.

### 4.2. Nanoplasmonic Assays: Functionalization and characterization

Gold nanoparticles (AuNPs) were synthesized using the standard citrate reduction method.^49^ This nano-plasmonic test was designed according to the published articles.^46,50–52^ Briefly, 2 mL of 50 mM HAuCl4 was added into 98 mL of boiling DI water in an Erlenmeyer flask. Then 10 mL of 38.8 mM sodium citrate was added, and the mixture was stirred until the color turned wine-red. The synthesized homogenous gold nanoparticles were characterized using UV-Vis spectroscopy and stored at 4°C. All aptamers were reconstituted in 1xPBS, 2mM MgCl_2_, pH 7.4 and targets were re-suspended in 1xPBS. All opioid aptamers were pre-heated at 95°C for 5 minutes to remove any dimerization before utilizing in any experiment. For aptamer validation, briefly, 100 nM of the aptamer is added to 100 μL of 13 nm sized AuNPs and incubated at room temperature (RT) for 5 mins. After 5 mins, 100 nM of target (n=NX, EDDP or NF) is added to the AuNP pre-incubated solution and wait additional 15-20 mins incubation at RT. Finally, ∼30 mM of NaCl is added to the final 100 µL solution. The nanoparticle-based color transition was observed immediately before and after adding the salt and recorded with a photograph. If the gold nanoparticle color is changed from red to purple, that indicates aptamer binding to the target. If the color does not change, then the target is not binding with aptamer. The change in the optical density (OD) at 520/700 nm (Abs_520/700_) of the resulting nanoparticle assembly was used to plot the aggregation rate and degree. UV-Vis’s spectra of each sample were measured in 96 well plates using BioTek microplate reader. Control experiments were performed in the absence of target (only Aptamer + AuNP + adjusted reaction buffer, 1xPBS, 2mM mgCl_2_, pH 7.4). We utilized a similar procedure for detection and validation of NX, EDDP, and NF in buffer and wastewater. In the previous study, the change in OD ratio at 520/700 nm of the resulting nanoprobe complex assembly was used to determine the limit of detection (3 σ/slope). ^46,50–52^

For sensitivity measurement, studies were also performed using various amounts (0, 25, 50, 100, 150 nM) of target NX in 100 μL of solution and, color changes were recorded immediately after incubation with ∼30 mM NaCl. Control experiments were performed in the absence of target NX. We performed similar procedure to detect opioids in wastewater where we spiked various amounts of target in presence of fixed aptamer concentration. The OD value at 520/700 nm and the pictures of the nanoparticle suspensions were recorded. All experiments were performed in triplicate (n=3) using 96 well plates. For specificity measurements, individual opioid aptamers with their target and/or non-target analyte with a ratio of aptamer: target=1:1 (for buffer) or 1:3 for WW was evaluated to verify its false positive and false negative binding performance. The change in OD value at 520/700 nm was measured and plotted. Similar experimental procedures were employed to other opioid targets (NF and EDDP) to perform the sensitivity and specificity measurements.

### 4.3. G-FET fabrication, Characterization, and Functionalization

#### 4.3.1. G-FET Fabrication and Characterization

G-FETs were fabricated with chemical vapor deposition (CVD) monolayer graphene transferred over SiO_2_/Si substrates. Monolayer graphene was grown on copper via low pressure chemical vapor deposition. The copper foil (Alfa Aesar) was pre-treated in Ni etchant (Transene) to remove any coatings or oxide layers from its surface. The tube furnace was evacuated to a read pressure of 200 mTorr with a constant flow of H_2_ (10 sccm). Prior to growth, the foil was annealed at 1010 °C (ramp rate 25 °C/min) for 35 minutes. Growth was done at 1010 °C with 68 sccm of H_2_ and 3.5 sccm of CH_4_ for 15 minutes. After growth, a polymethyl methacrylate (PMMA) layer was spin coated on one side of the copper foil and baked for 60 seconds at 60 °C. To facilitate smooth and fast etching of the copper foil, the backside graphene was etched using oxygen plasma with 60 Watt power for 60 seconds. The exposed copper was etched away in Nickel etchant for 2h at 60 _o_C. The remaining PMMA/graphene structure was washed in three DI water baths, the first and second water baths for 60 seconds each and the third for 30 minutes, to rinse away left-over etchant. The source/drain along with coplanar gate electrodes were patterned on SiO_2_/Si chips of size 1.2 cm × 1.2 cm using bilayer photoresist (LOR1A/S1805) and laser mask writer (Heidelberg Instruments) followed by Pt/Ti (20 nm/5 nm) deposition with e-beam (Angstrom Engineering) and lift off using remover PG (MicroChem). To remove photoresist residues and improve the adhesion of electrodes, a 10 h baking was performed at 400 °C in vacuum. The PMMA/graphene was then transferred onto these prepatterned Pt/Ti electrodes. Any leftover water was slowly dried with argon gas, and finally the PMMA was dissolved in acetone vapors; IPA (Fisher) was used for a final wash. The chips were baked at 300 °C for 8h in vacuum to ensure graphene adhesion and further clean photoresist residue. This was followed by deposition of 3 nm AlO_x_ at room temperature by e-beam deposition to protect the graphene. Substrates were baked at 175 °C for 10 minutes before lithography process. After that the graphene patterning was done with lithography using same bilayer resists and then etched with oxygen plasma for 30s at 75 Watt followed by 3 minutes of Argon plasma at 100 Watt to remove any oxide formed over Pt gate electrodes. Devices were cleaned with remover PG and rinsed with IPA, DI water and dried with Argon followed by removal of the 3 nm AlO_x_ layer by dipping in MF-321 developer for 30 seconds. Then, for electrode passivation to protect the electrodes and edges of the graphene for liquid gating, 50 nm AlO_x_ was deposited using e-beam and AlOx crystals (Lesker) at oxygen pressure of 7.5 × 10^5^ mbar. Photolithography was done using S1805 to expose the sensing area (10 × 40 µm), gate electrodes, and contact pads while leaving remaining chip covered. The chips were post baked at 120 °C for 5 minutes followed by AlO_x_ etching in transetch (Transene) for 7:30 minutes at 80 °C hot plate temperature. To hold the solution for experimental measurements and for functionalization, PDMS wells of size 1.5 × 1.2 mm were fabricated and placed over the chip segregating the four sets of devices with five devices in each well.

#### 4.3.2. G-FET Functionalization

Fabricated G-FETs were functionalized with respective aptamers for specific and selective detection of opioids. G-FET chips were incubated for 1h with high concentration (10 mM) PBASE linker dissolved in DMF. Next, the G-FET was rinsed with DMF to remove adsorbed linker molecules followed by rinsing with IPA, DI to clean the surface of solvents. The pyrene group in PBASE linker stacks over the graphene surface through *π-π* interaction while the N-hydroxysuccinimide (NHS) ester reacts with amine terminated at 5’ end of aptamers _23_. Chips functionalized with linker were incubated for 1h with PBS solution with 2mM MgCl_2_ containing aptamers with a concentration of 10 μM. Finally, the chips were rinsed with PBS to remove excess aptamers, followed by DI to clean the salts from the graphene surface.

## ASSOCIATED CONTENTS

### Supporting Information

Raman spectra of graphene before and after fabrication. G-FET characteristics obtained from a chip having all twenty devices with 4 different PDMS wells. NF and EDDP detection results in wastewater. EIS based detection results of NX with sensitivity and selectivity. Raman spectra to confirm the functionalization. G-FET characteristics for detection of NX in 1x PBS and in wastewater. Calibration curve for NX detection in wastewater without using PEG.

## AUTHOR INFORMATION

### Corresponding Authors

*

*

### Author contributions

K.S.B., A.A., N.K., M.R. conceived the project and designed the experiments. N.K., performed fabrication, functionalization, and testing of G-FETs. M.R, N.I.K, performed the aptamers validation with nanoplasmonic and electrochemical assays, respectively. M.R., A.W., B.D. selected the aptamers and opioid metabolites. M.G. assisted in the fabrication and characterization of G-FETs. M.C. developed the PCB circuit board, mold, and optimized the PDMS wells, J.O-M. helped with functionalization of devices and selectivity tests, K.H. performed the growth of graphene on copper and helped in transferring over to SiO_2_/Si substrates, X.L. supervised the graphene growth and transfer process. N.K, M.R., K.S.B, A.A., M.G., N.I.K., T.v.O analyzed the data and wrote the manuscript.

^#^ These authors contributed equally.

## Acknowledgements

N.K. is grateful for the support of the for the support of the Office of Naval Research under Award number N00014-20-1-2308. The work of M.G., M.C., and K.S.B. was supported by the National Science foundation via award DMR-2003343. M.R, N.I.K, A.W., B.D, and A.A acknowledge the funding support from National Institutes of Health National Institute on Drug Abuse (NIDA) through SBIR Phase I grant (Award # 1R43DA051105-01). J.O-M and T.v.O were supported by the National Institutes of Health, through grant U01-AI124302. X.L. and H.K. acknowledge the support of the National Science Foundation (NSF) under Grant No. (1945364). X.L. acknowledges the membership of the Boston University Photonics Center. We would like to thank Mr. George Heufelder, Director of MASSTC for providing us wastewater influent samples for this study and Prof. Kurunthachalam Kannan at Department of Environmental Medicine, NYU for helpful discussions and guidance.

