## Supporting Information for "Rapid, multianalyte detection of opioid metabolites in wastewater"


### SI 1.1. GFETs' fabrication and characterization

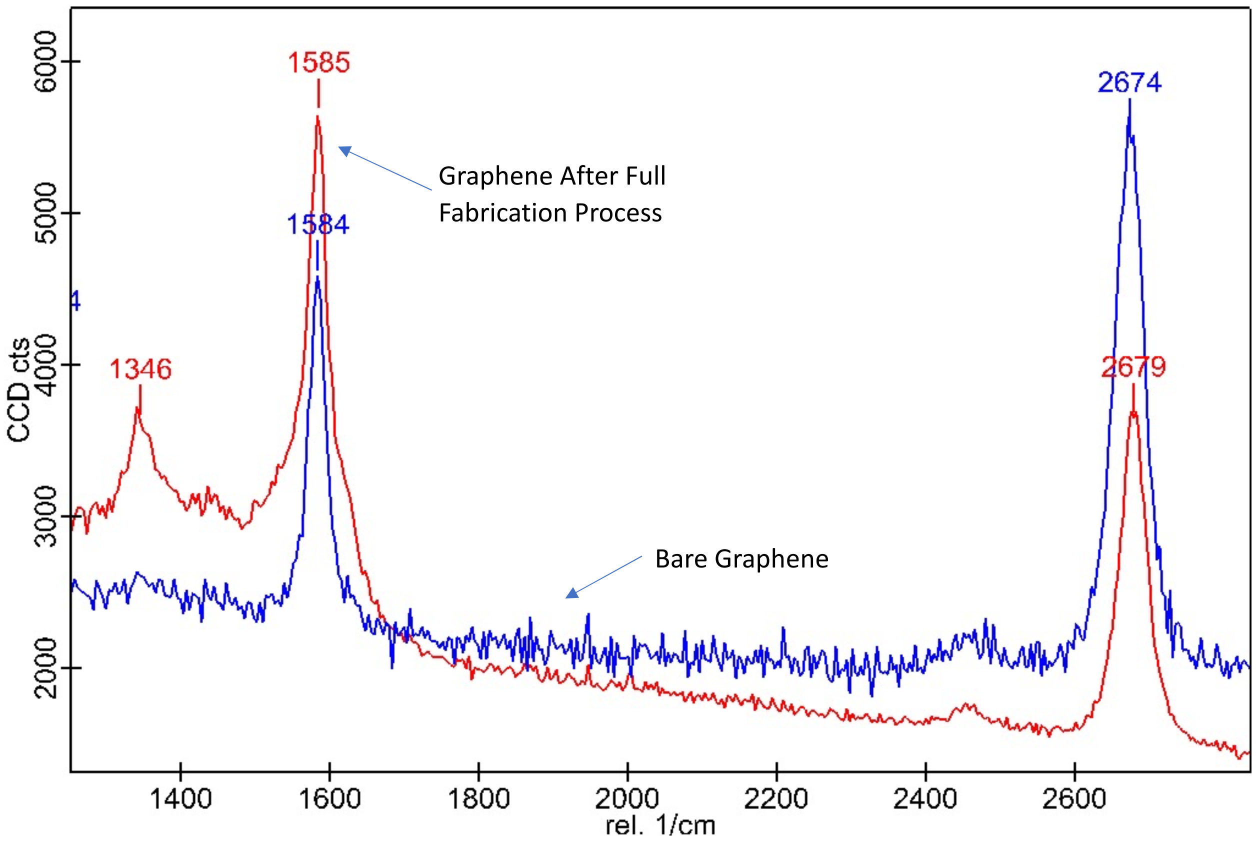

**Figure S1.** Raman spectra showing D, G and 2D peaks before and after fabrication process. The G peak shows little shift from 1584 $cm^{-1}$ to 1584 $cm^{-1}$ signifying negligible doping of the graphene occurred during the fabrication process. The reduction in 2D peak is due to non-charge carrying contaminants/defects introduced during fabrication. This could arise from chemicals and possible contaminants on graphene, electrodes, and Si/SiO2 substrate. The increase in D peak implies some additional, charge neutral disorder added during fabrication.

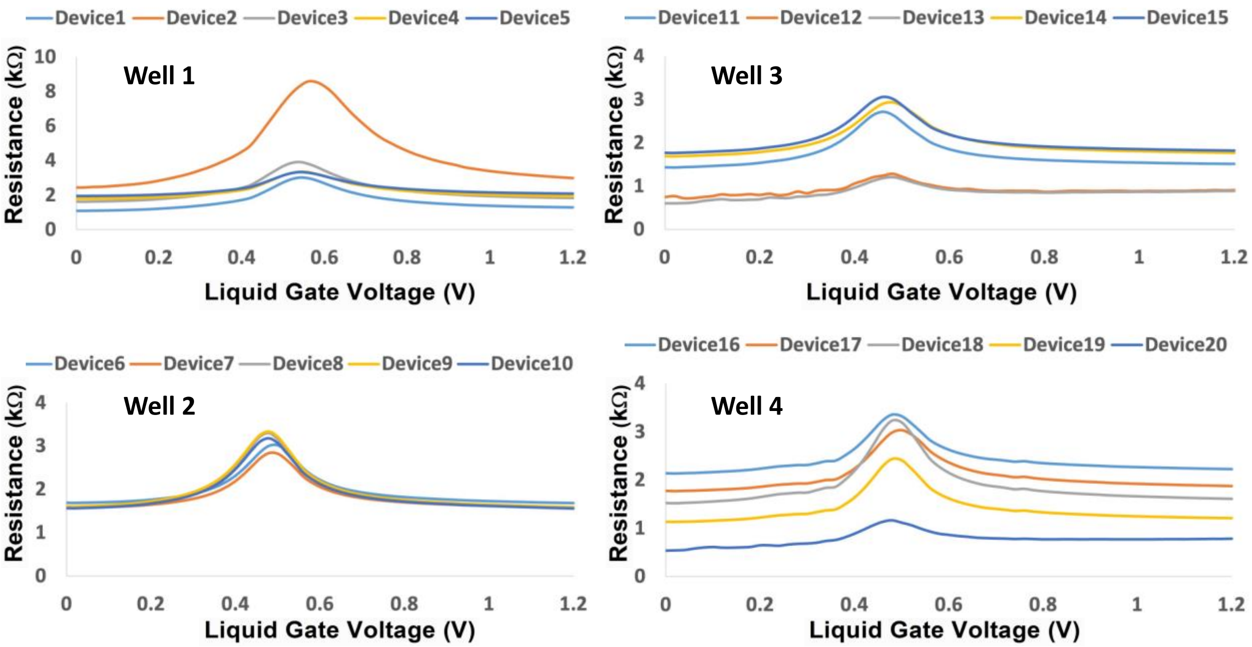


**Figure S2.** Initial Dirac Voltage ($V_D$) of each well occurs at peak resistance. All data are from the four separate wells of a single AptG-FET chip showing all twenty devices working. The initial $V_D$ for each device is around 0.5 V. This non-zero value of $V_D$ is due to the higher electrochemical potential of the Platinum electrode than the typical reference electrode (i.e. Ag/AgCl). The higher resistance for Well 1 – Device 2 may be due in part to patches in graphene, unetched AlOx present in graphene window, or presence of some contaminant/defect.

#### SI 1.2. Detection of opioid metabolites in wastewater using nanoplasmonic assay

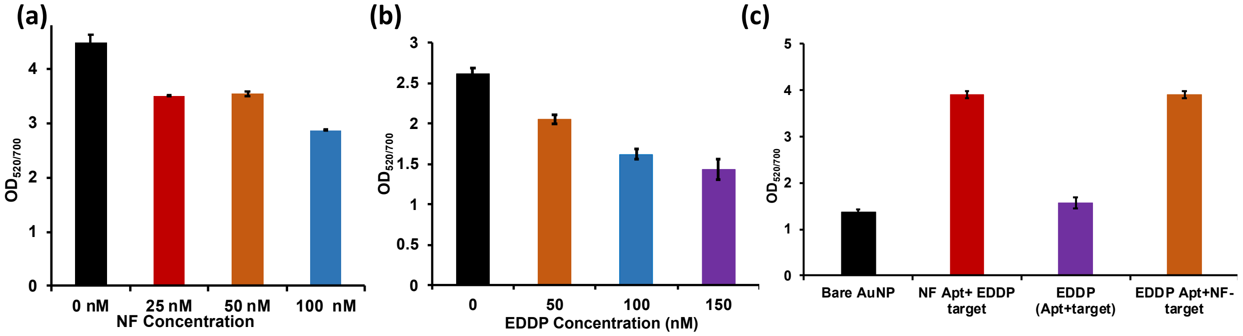


**Figure S3.** Characterization of an Opioid aptamer (such as NF and EDDP) that includes sensitivity (a-b) in buffer and specificity (c) analysis in wastewater samples. All experiments were performed in triplicate (n=3).

### SI 1.3. Electrochemical impedance spectroscopy (EIS): Detection methodology

#### Experimental Details

For characterizing the sensor, the electrochemical impedance spectroscopy (EIS) measurements were performed at each modification step. EIS measurements were conducted in the presence of 1mM $K_4[Fe(CN)_6]/K_3[Fe(CN)_6]$ (1:1) mixture (pH 7.4) as a redox probe in 1x PBS (phosphate buffer saline (PBS)). The impedances were measured over a frequency range from 0.5 Hz to 50 kHz at a DC offset of 120 mV versus the open-circuit voltage (OCP) with a sinusoidal signal of 10 mV RMS. The sampling rate was 10 points per decade. After the EIS measurements, each Nyquist curve was fitted using the Randel's circuit to obtain the charge transfer resistance ($R_{ct}$) as the sensing parameter.

Each measurement was done with a 50 µL drop of corresponding redox probe solution placed on the electrode followed by a rinsing step with PBS buffer. First, without any modification on the screen-printed carbon electrode (SPCE), 50 µL redox probe was placed on the electrode and the EIS measurements were performed for bare condition. Then each SPCE was incubated in 40 µL of 1 µM aminated aptamers suspended in 0.01x PBS containing 1 mM $MgCl_2$ for aptamer immobilization using the covalent coupling between the carboxyl groups attached to the SPCE surface and the amino groups attached to the 5′ end of the aptamers. Each electrode was then rinsed with the PBS buffer to remove the unbound aptamers from the electrode surface and EIS measurements were taken by placing 50 µL of redox probe on the aptamer modified SPCE. Next, each device was exposed to 40 µL of NX of varying concentrations (0, 10, 100, 1000 nM) and incubated for 1 hour for target binding. Following the target incubation, each electrode was rinsed with PBS buffer to remove any unbound NX. Afterwards, 50 µL of redox probe was dropped onto the electrode surface and EIS measurements were performed again.

### Results

To further confirm the aptamers binding for NX detection, we also performed electrochemical impedance spectroscopy (EIS) using commercially available screen-printed electrodes. EIS is an excellent and sensitive tool for monitoring any change at the electrolyte and electrode interface and very commonly used for characterizing the features of surface-modification (Figure S4).[1] Figure S4 shows an absence test for confirming the specific binding of the aptamer-NX using the EIS. As can be seen in Figure S4a, with the bare screen-printed carbon electrode (SPCE) exposed to 100 nM of NX, there is no change in the charge transfer resistance ($R_{ct}$), while upon anchoring of aptamers on the electrode, the Nyquist curve jumps up leading to an increase in the electron transfer resistance. This increase in resistance can be explained by the negative charge of the DNA backbone. After aptamer functionalization, the modified electrode was incubated in 1x PBS having zero concentration of NX. Although, no $R_{ct}$ change is expected for this case, the reduction (~8%) in the $R_{ct}$ is possibly due to the removal of some functionalized aptamers from the electrode surface in the incubation step. However, upon exposure of 100 nM NX, the charge transfer resistance increases again. This consistent increase in the resistance is due to the increased concentration of bound NX that hinders the charge transfer**.** In Figure S4b, in presence of various concentration of NX target with NX aptamer we observed a dose-dependent curve. Figure S4c shows that when high concentration of non-target is introduced, no increase in the electron transfer resistance is observed, which confirms the aptamer binding specificity with no false positive/negative interference.

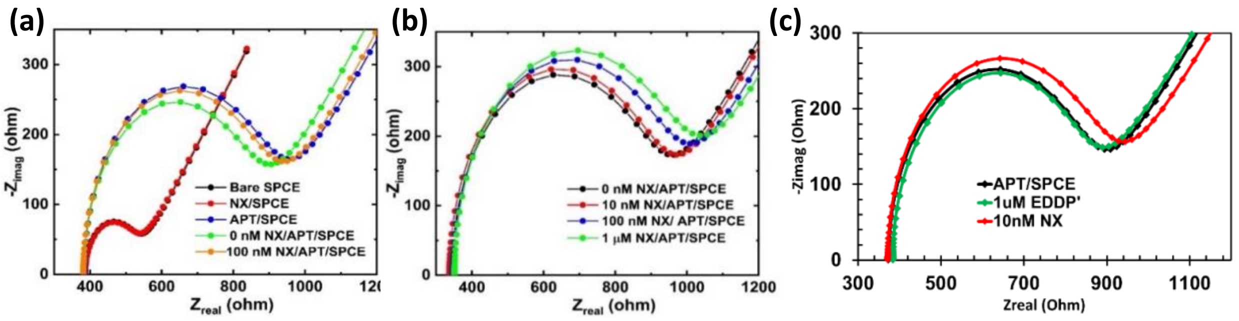


**Figure S4.** EIS for characterizing aptamer-NX binding: (a) EIS showing the specific affinity of the aptamer to NX target, and (b) EIS showing the dose-dependent response of the sensor to NX target, (c) Specificity testing using NX aptamer with NX target and/or unspecific target (EDDP).

**SI 1.4. G-FETs' functionalization and detection of opioid metabolites**

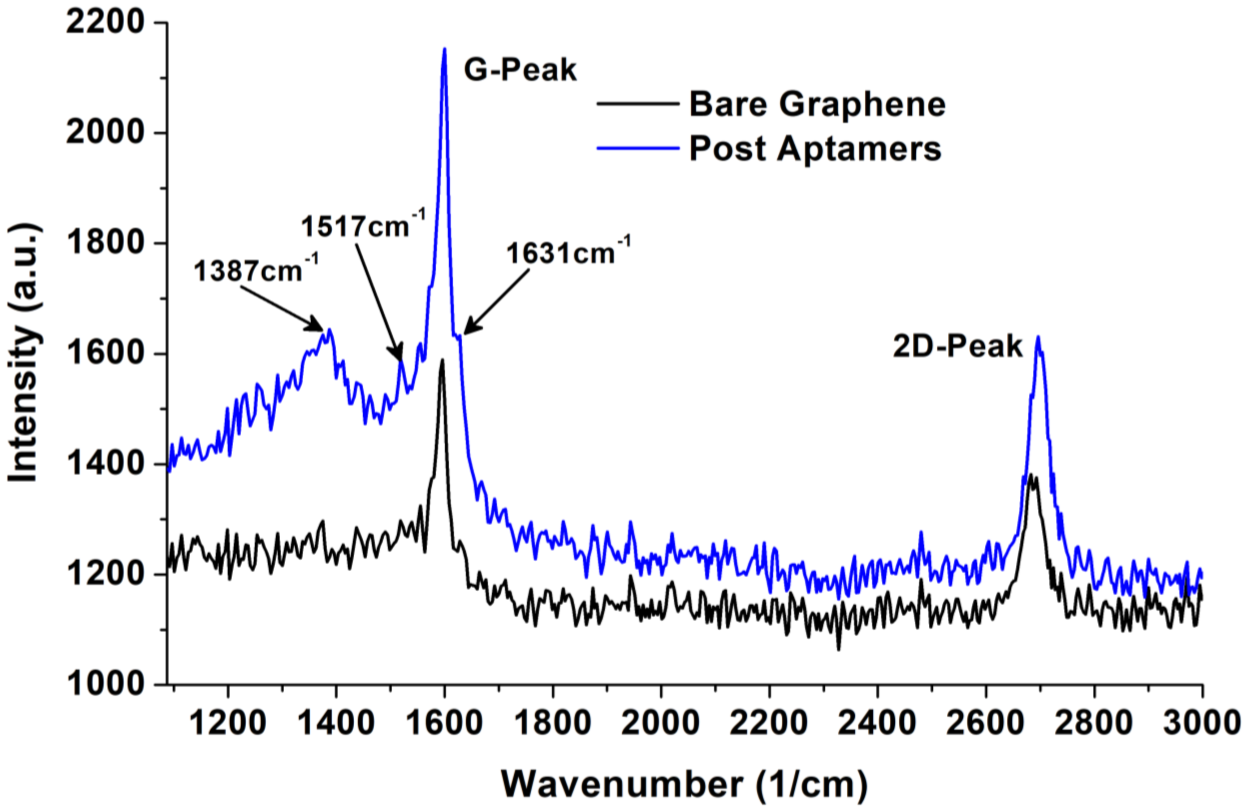


**Figure S5.** Raman spectra of graphene over $SiO_2/Si$ substrates before and after aptamers' functionalization, emergence of three new peaks at 1387, 1517, and 1631cm$^{-1}$ confirmed the attachments of aptamers over graphene

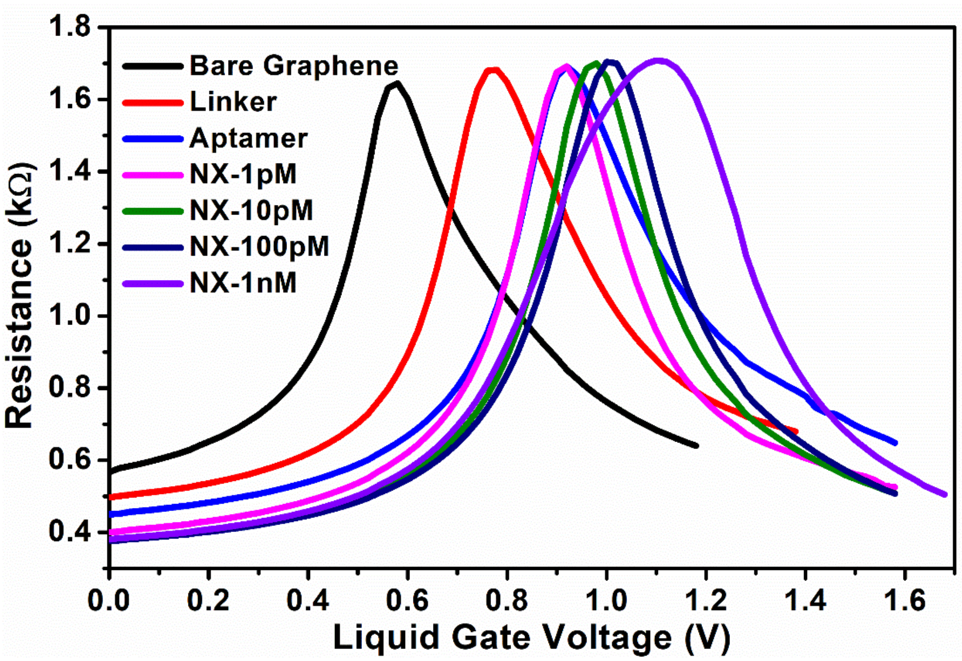


**Figure S6.** Detection of NX in 1x PBS with successive increase in concentration

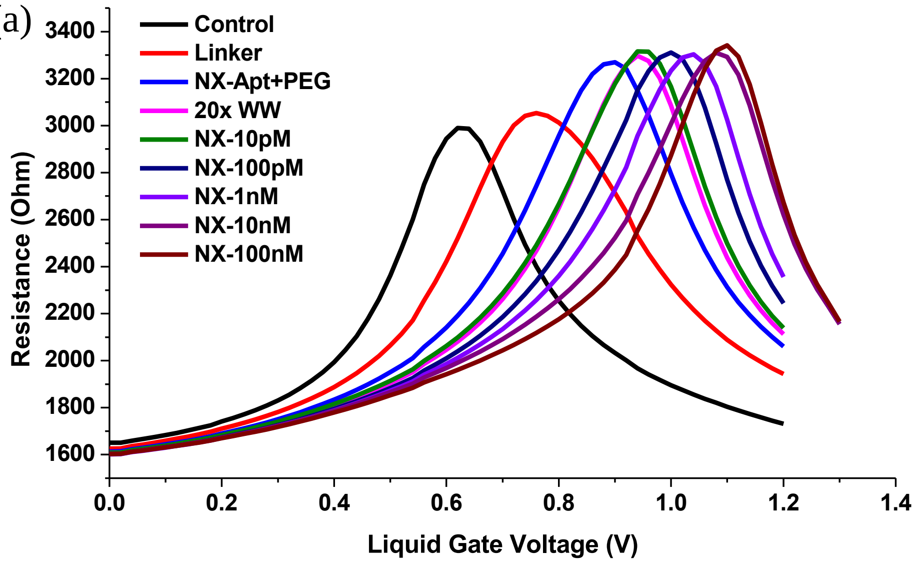

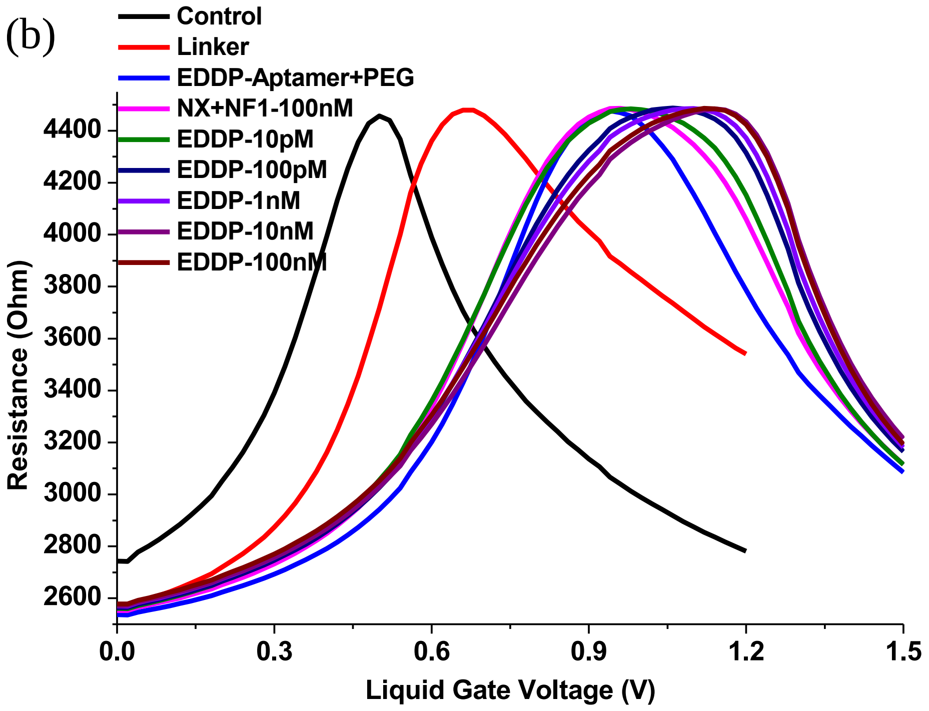


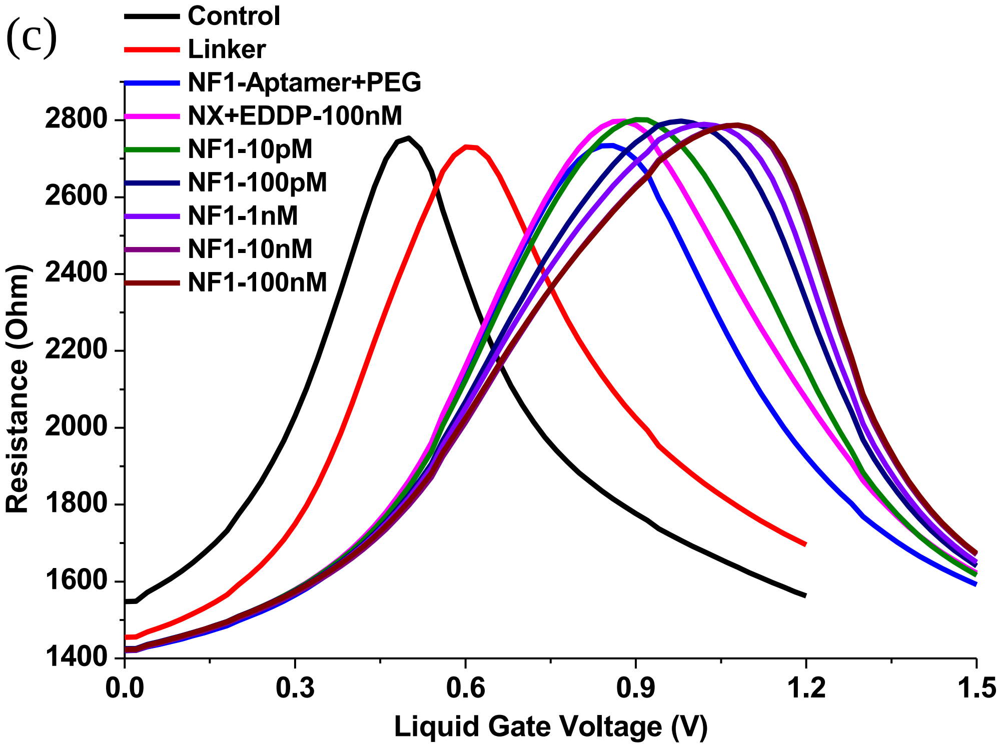


**Figure S7.** Representative Resistance vs Voltages plots of G-FET devices at each stage of functionalization and different concentrations of NX, EDDP, and NF

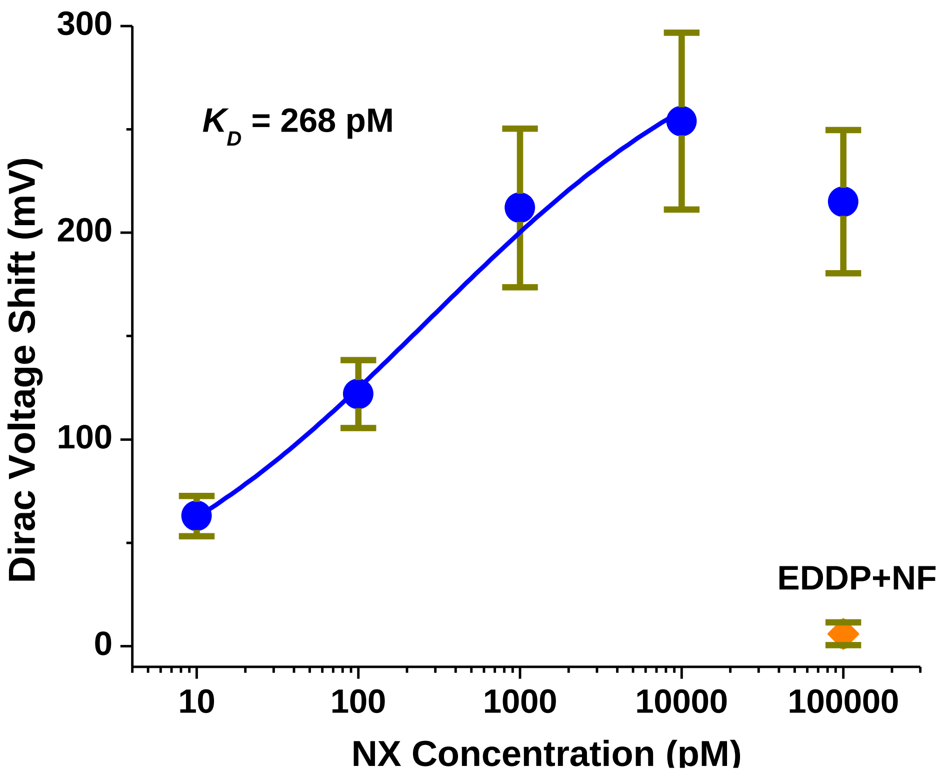


**Figure S8.** AptG-FET without PEG, voltage shift with different concentration of NX target
